# BARTweb: a web server for transcription factor association analysis

**DOI:** 10.1101/2020.02.17.952838

**Authors:** Wenjing Ma, Zhenjia Wang, Yifan Zhang, Neal E. Magee, Yang Chen, Chongzhi Zang

## Abstract

Identifying active transcription factors (TFs) that bind to cis-regulatory regions in the genome to regulate differential gene expression is a key task in gene regulation research. TF binding profiles from numerous existing ChIP-seq data can be utilized for association analysis with query data for TF identification, as alternative to DNA sequence motif analysis. Here, we present BARTweb, an interactive webserver for identifying TFs whose genomic binding patterns associate with input genomic features, by leveraging over 13,000 public ChIP-seq datasets for human and mouse. Using an updated Binding Analysis for Regulation of Transcription (BART) algorithm, BARTweb can identify functional TFs that regulate a gene set, or have a binding profile correlated with a ChIP-seq profile or enriched in a genomic region set, without a priori information of the cell type. Compared with the original BART package, BARTweb substantially reduces the execution time of a typical job by two orders of magnitude. We also show that BARTweb outperforms other existing tools in identifying true TFs from collected experimental data. BARTweb is a useful webserver for performing functional analysis of gene regulation. BARTweb is freely available at http://bartweb.org.

## INTRODUCTION

Transcription factors (TFs) play an instrumental role in controlling gene expression by interacting with regulatory DNA elements in the eukaryotic genome (1). An important task in gene regulation studies is to identify active TFs that function to regulate genes with differential expression or are enriched in certain regions in the genome. Chromatin immunoprecipitation followed by high-throughput sequencing (ChIP-seq) has become one of the most commonly used techniques for genome-wide profiling of TF binding sites and chromatin marks (2, 3). The increasingly large amount of publicly available ChIP-seq datasets generated by individual laboratories worldwide as well as large collaborating consortia such as Encyclopedia of DNA Elements (ENCODE) (4) and Roadmap Epigenomics (5) is a valuable resource for interrogating genomic profiles for hundreds of TFs in many human and mouse cell types (6). As an alternative to DNA-binding sequence motif search, compendium ChIP-seq data collected from the public domain can be utilized to perform TF identification or association analysis, as ChIP-seq carry direct binding information of TFs and cover the entire genome, most of which is non-coding but contains regulatory elements such as enhancers.

To leverage publicly available ChIP-seq data for TF identification, we previously developed Binding Analysis for Regulation of Transcription (BART), an algorithm to identify TFs from a large collection of ChIP-seq data that have a genomic binding pattern highly correlated with an input genomic profile, using a novel statistical approach integrating multiple levels of statistical tests (7). To predict TFs regulating a query gene set, BART first apply Model-based Analysis of Regulation of Gene Expression (MARGE) (8) to derive a genomic cis-regulatory (enhancer) profile from the input gene set using a semi-supervised learning approach leveraging compendium ChIP-seq data for active enhancer histone mark H3K27ac, then generate a ranked list of TFs that have a highly correlated binding profile with the cis-regulatory profile. While proven to work for identifying functional TFs from many case studies (7, 9–12), BART requires users to download large ChIP-seq data libraries that can be storage and memory-consuming, and sometimes runs slow primarily due to step-wise regression computation in MARGE.

There are several other bioinformatics tools that also use existing ChIP-seq data for TF identification or enrichment analysis, such as TFEA.ChIP (13) and ChEA3 (14). TFEA.ChIP applies the Fisher’s exact test or the Gene Set Enrichment Analysis (GSEA) method (15) for TF enrichment analysis using ChIP-seq data collected from the ReMap database (16). ChEA3 integrates multiple sources of TF-target association information including ChIP-seq, co-expression from RNA-seq, and collected crowd-based gene lists (17) to generate a ranked list of TFs associated with query gene sets. HOMER (18) and Pscan (19), representing conventional sequence motif-based methods for TF identification from target gene sets, are also included in our work for comparison and evaluation purposes.

To overcome several data-intensive computing burdens and to improve the performance of the original BART package, we present BARTweb, a web-server application for users to perform transcription factor identification analysis associated with multiple types of query data. BARTweb is accessible through a user-friendly web interface, from which users can submit three types of input data: a gene set, a ChIP-seq mapped read dataset, or a scored genomic region set. The output includes a table of TFs ranked by a series of quantification scores with statistical assessments, as well as several analysis plots for each TF. BARTweb incorporates an updated data library, which includes over 13,000 high-quality ChIP-seq datasets for over 900 human transcription factors and over 500 mouse transcription factors. BARTweb also implements an updated BART algorithm applying adaptive Lasso (20) in place of MARGE step-wise regression for faster and more robust performances. We demonstrate that BARTweb outperforms several existing tools in identifying true TFs from curated experimental data, and can be a useful tool for the gene regulation research community.

## MATERIAL AND METHODS

### BARTweb server infrastructure design

In order to provide a user-friendly and stable service through web interface, we designed a two-part structure for the BARTweb server: a front-end web interface part and the back-end computing service (Figure 1). The front-end layer is used to receive users’ job submission requests and to display job execution information and results. The back-end service performs all of the computation. We containerized both parts into Docker and deployed them on a 17-server Distributed Cloud Operating System (DCOS) cluster for continuous and stable services.

**Figure 1.**
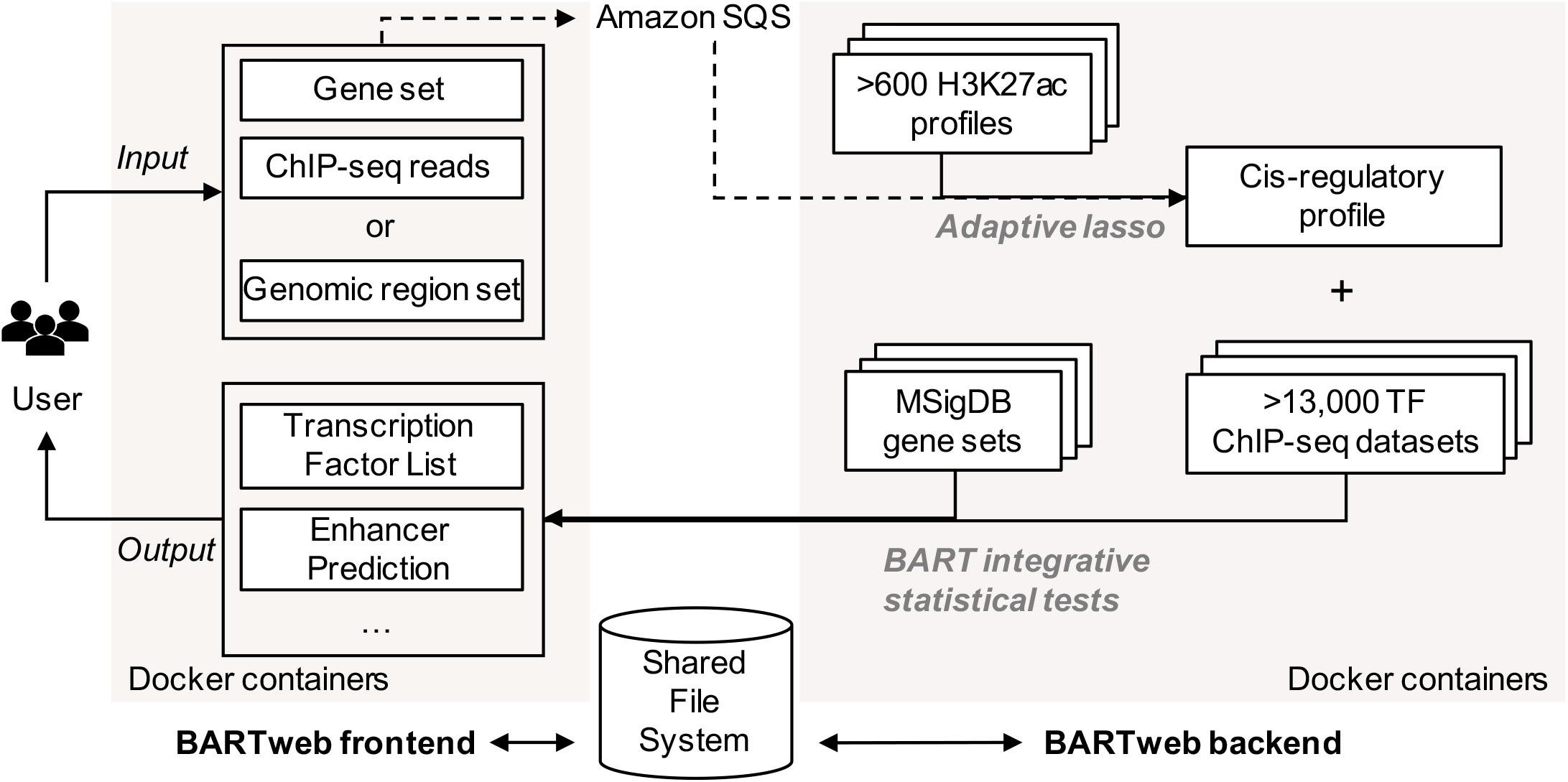
BARTweb architecture overview. BARTweb front-end receives user input and displays processed output. BARTweb back-end performs computation of BART TF identification analysis. Both services are containerized and share a common file system.

The front-end web interface was developed and implemented in Flask, a lightweight, easy-to-extend, Python syntax-based framework. To support simultaneous users, we deployed it under Apache 2.4 inside a Docker container. The back-end service uses our updated BART algorithm implemented in Python3. To ensure continuous deployment of the website, we serialized both parts into a Github repository and use Travis Continuous Integration (Travis-CI) to automatically push the code changes into the online environment running in production.

To connect the front-end and the back-end, as well as to allow the application to scale to many users, we employed a robust queue using Amazon’s Simple Queue Service (SQS) to temporarily store job keys. Every time a user submits a new job request through the web interface, the BARTweb front-end pushes a message unique to that request into the queue. Meanwhile, the BARTweb back-end routinely checks that queue for incoming requests, executes as soon as a new job comes in, and removes the request from the queue. Using a message queue also ensures that if the back-end service is unavailable, the messages submitted by users are still captured and can be processed later.

We generate a unique key and a corresponding uniform resource locator (URL) once a job is submitted. Users can leave BARTweb running in the background or store the key for later retrieving the results. Users have the option to provide their email addresses for receiving notifications about the job status and the result URL. The results are kept on the server with the unique keys or URLs for a minimum of 180 days, so that users can retrieve and share the results anytime for a while after running the job. BARTweb does not store user submitted data.

### Updated BART algorithm

In BARTweb, we applied an updated BART algorithm, in which the inference of the genomic cis-regulatory profile from the input gene set by integrating compendium H3K27ac ChIP-seq data was replaced from the original MARGE algorithm (8). MARGE adopts a forward stepwise regression for feature selection to identify significant predictors, i.e., informative H3K27ac profiles that carry regulatory potential information to better separate the input gene set from other genes in the genome. However, stepwise regression has fundamental limitations including selecting extremely variable features and frequently trapped into a local optimal solution. In addition, k-fold cross-validation makes the entire job execution very slow. In the updated algorithm, we use adaptive lasso (20) for this feature selection process.

Similar to MARGE, we consider the selection of informative H3K27ac samples as a logistic regression model. Suppose ***y*** = (*y*_1_, … *y*_*n*_)^*T*^ be the response vector indicating whether a gene belongs to a given gene set or not, and ***P*** = [***p***_1_, … ***p***_*m*_] be the predictor matrix, i.e., the normalized regulatory potential (RP) matrix derived from H3K27ac profiles (8). We assume that

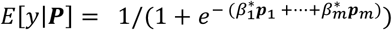

We further assume 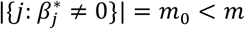, and the model to estimate the input gene set depends only on a sparse representation of the predictors, i.e., a small subset of samples from the H3K27ac ChIP-seq data compendium. We use adaptive lasso to identify an accurate sparse representation of the predictors. The generalized logistic adaptive lasso is defined as

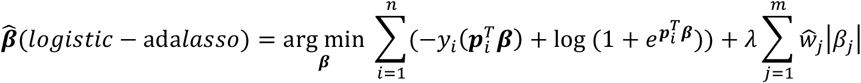

The adaptive lasso carries the oracle properties, namely, it can simultaneously achieve consistent variable selection and optimal prediction rate. Compared to lasso, which equally penalizes the coefficients in the ℓ_1_ penalty, adaptive lasso uses data-dependent adaptive weights to penalize different coefficients in the ℓ_1_ penalty. The weight vector can be selected based on the importance of different indictors, e.g., large coefficients do not be penalized much as the corresponding variables can be important, and the irrelevant variables are penalized more. By performing a different regularization for each coefficient, the adaptive lasso avoids over-penalization of relevant coefficients, reduces the estimation biases, and leads to a consistent model selection (21). Besides, adaptive lasso is asymptotically as efficient as the least squares regression using a proper model. By applying the LARS algorithm (22) that is implemented in our model, the adaptive lasso is in the same order of computation of a single ordinary least squares (OLS) fit (20).

If the weights are cleverly chosen, the adaptive lasso performs equally well as if the true underlying model were given in advance (20). The weights can be initiated as 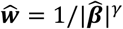, where 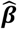 can be the least-squares estimator 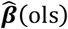 or the ridge regression estimator 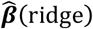, and *γ* > 0. Here we iteratively construct the data dependent adaptive weights. The weights are initiated as an all-one-vector, and then are iteratively determined by the coefficient from the logistic lasso in the previous step. The algorithm is described in Supplementary Data: Algorithm 1.

After relevant H3K27ac samples are selected, we directly apply the feature coefficients on the H3K27ac signals for each candidate enhancer site (union DNase hypersensitive site, UDHS) and produce a score as the enhancer prediction, instead of using the K-means-based semi-supervised learning in MARGE. The higher the score is, the more likely this site is a functional element regulating the input gene set. All candidate enhancer sites with prediction scores compose the genomic regulatory profile, which undergoes the rest steps in the BART algorithm.

### Updated ChIP-seq data library

The amount of available ChIP-seq data keeps growing in the public domain. In order to increase the analysis power, we updated the ChIP-seq data library to cover more TFs in more cell types for both human and mouse. We downloaded the TF ChIP-seq peak files from the updated version of Cistrome Data Browser (23). Under the same quality control standards used in BART v1.1 (7), we processed the peak data and kept only the datasets that have at least 2,000 peaks. The current data library contains 7,968 ChIP-seq datasets for 918 human TFs and 5,851 ChIP-seq datasets for 565 mouse TFs, a significant increase from BART 1.1 (Figure 2A, B). It is worth noting that although histone modification ChIP-seq data were excluded, we still keep many chromatin regulators (e.g., EZH2) and some histone variants (e.g., H2A.Z) in the TF ChIP-seq data library. Although these chromatin factors do not necessarily bind DNA directly and usually are not categorized as “transcription factors”, many of them still have gene regulation-related functions. Therefore, we still include them to maintain a diverse collection of regulatory factors and a strong analysis power. For simplicity in terminologies, we refer to all DNA-associating protein factors as transcription factors (TFs), without detailed separation between DNA-binding TFs and chromatin regulators, etc.

**Figure 2.**
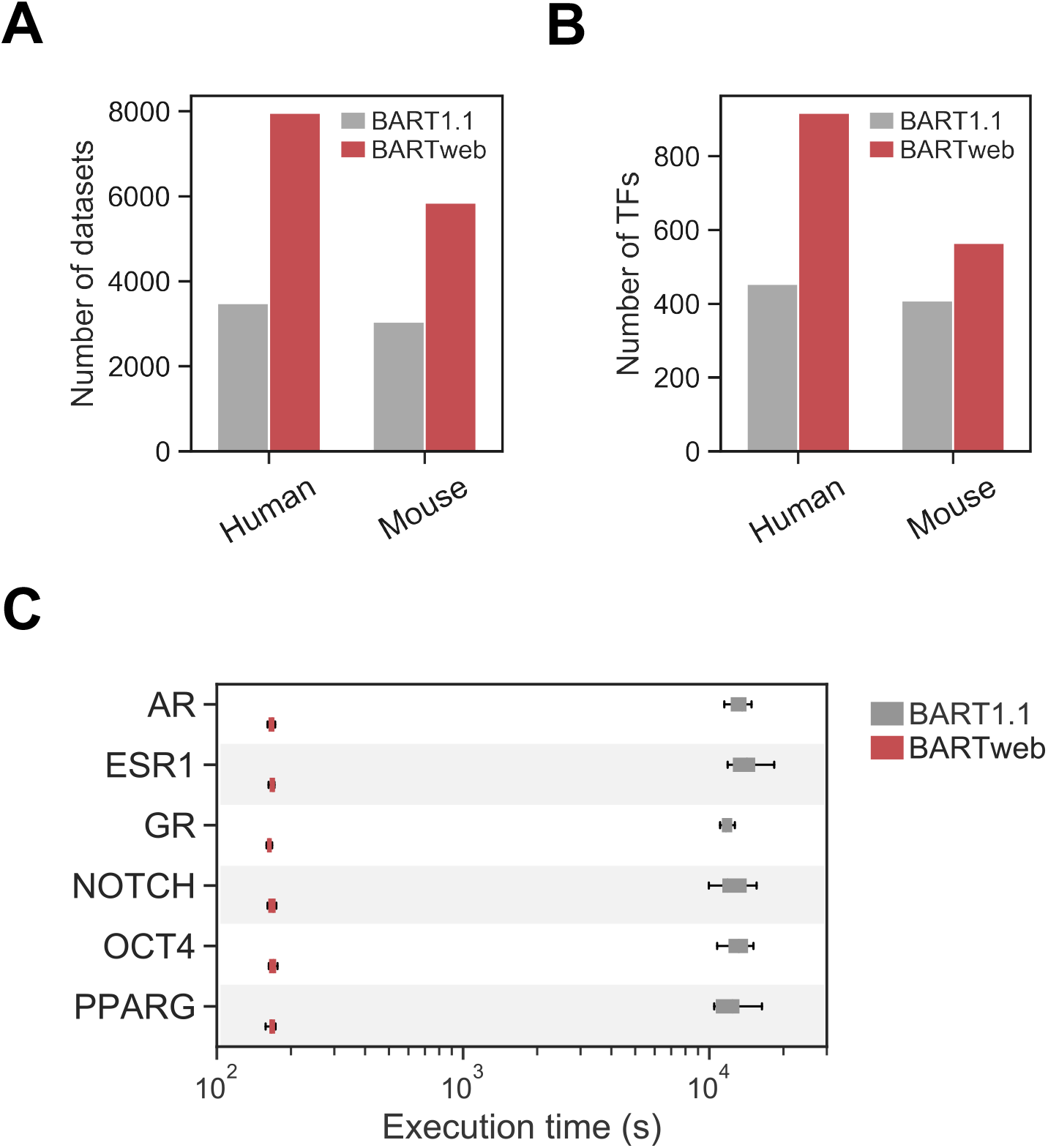
BARTweb improvements over BART1.1. (**A,B**) Bar plots showing the number of collected ChIP-seq datasets (**A**) and the number of uniquely covered TFs (**B**) in BARTweb and BART1.1 data libraries. (**C**) Box plots showing execution times on analysing six differentially expressed gene sets, comparing between BARTweb and BART1.1. Each gene set was run 20 times.

Other data in the library largely remain the same as BART 1.1, including UDHS data sets with over 2.7 million sites covering 307Mbp regions in the human genome and over 1.5 million sites covering 121Mbp regions in the mouse genome. For background models, we collected 514 gene sets with over 200 genes from the Chemical and Genetic Perturbation sets of Molecular Signature Database (MSigDB) (24) for human gene set input, 366 H3K27ac ChIP-seq profiles for human ChIP-seq input and 267 H3K27ac profiles for mouse input. For each dataset, we pre-calculated the Wilcoxon statistic scores with the new BART algorithm and generated the background standardization matrix. We plan to keep updating the data library on a semi-annual basis.

We also updated the data structure of storing each TF occupancy information in UDHS to improve both storage efficiency and execution time. The original TF binding occupancy matrices, where each row represents a UDHS and each column a TF dataset, are extremely sparse matrices, in which only 1.39% or 2.05% elements are occupied by true TF binding, for human and mouse, respectively. Therefore, we only stored the true binding TF dataset information for each UDHS, and discarded other information. In this way, we could save storage space and increase running speed at the same time.

## RESULTS

### BARTweb improvements on TF coverage and running time

We demonstrated the performance of BARTweb using the six public differentially expressed gene sets upon different TF perturbation that were used in BART (7). We found that with updated data library (Figure 2A, B) and improved algorithm, the running time for BARTweb was substantially reduced compared with BART v1.1, from nearly 3 hours to less than 5 minutes (Figure 2C), for all 6 TFs. Meanwhile, the prediction accuracy remains high (Supplementary Figure S1).

### BARTweb input

BARTweb can accept any one of the three data types as input:

i. *a gene set* in official gene symbols in text format. BARTweb will first integrate the gene set with H3K27ac ChIP-seq data compendium to derive a genomic cis-regulatory profile, which represents the ranked list of functional regulatory elements (enhancers) that are predicted as more likely to regulate the input gene set compared to other genes in the genome. Then TF association analysis was performed on this genomic cis-regulatory profile. At least 100 genes are recommended in the input.
ii. *a ChIP-seq mapped read dataset* in BAM or BED format. BARTweb will first pile up the ChIP-seq reads located at the UDHS, and use the read count at each UDHS site to generate a ranked list of candidate regulatory elements, or the genomic regulatory profile, to perform TF association analysis. At least 1 million reads are recommended in the input.
iii. *a scored genomic region set* in 4-6 column BED format. BARTweb will first map the region set to UDHS, and assign the region score to each UDHS overlapped with the region to generate the ranked list of candidate regulatory elements, or the genomic regulatory profile, to perform TF association analysis. At least 1,000 regions are recommended in the input.

Currently, only human (hg38) or mouse (mm10) are supported for mapped ChIP-seq read data or region set annotations. When submitting a job through the web interface, users need to specify the species and input data type besides providing the input data. The input data can be either uploaded as a file in a correct format, or pasted in the input field.

### BARTweb output

Once a job is submitted, a status indicator and a processing log will be displayed on the web interface. BARTweb usually takes around 3-5 minutes to run the job, then it will display the result panel, including a ranked list of all TFs with quantification scores (Figure 3A) and a list of all intermediate or final output data files available for download. For each TF, clicking on the TF name will open a pop-up window displaying its corresponding analysis plots (Figure 3B, C).

**Figure 3.**
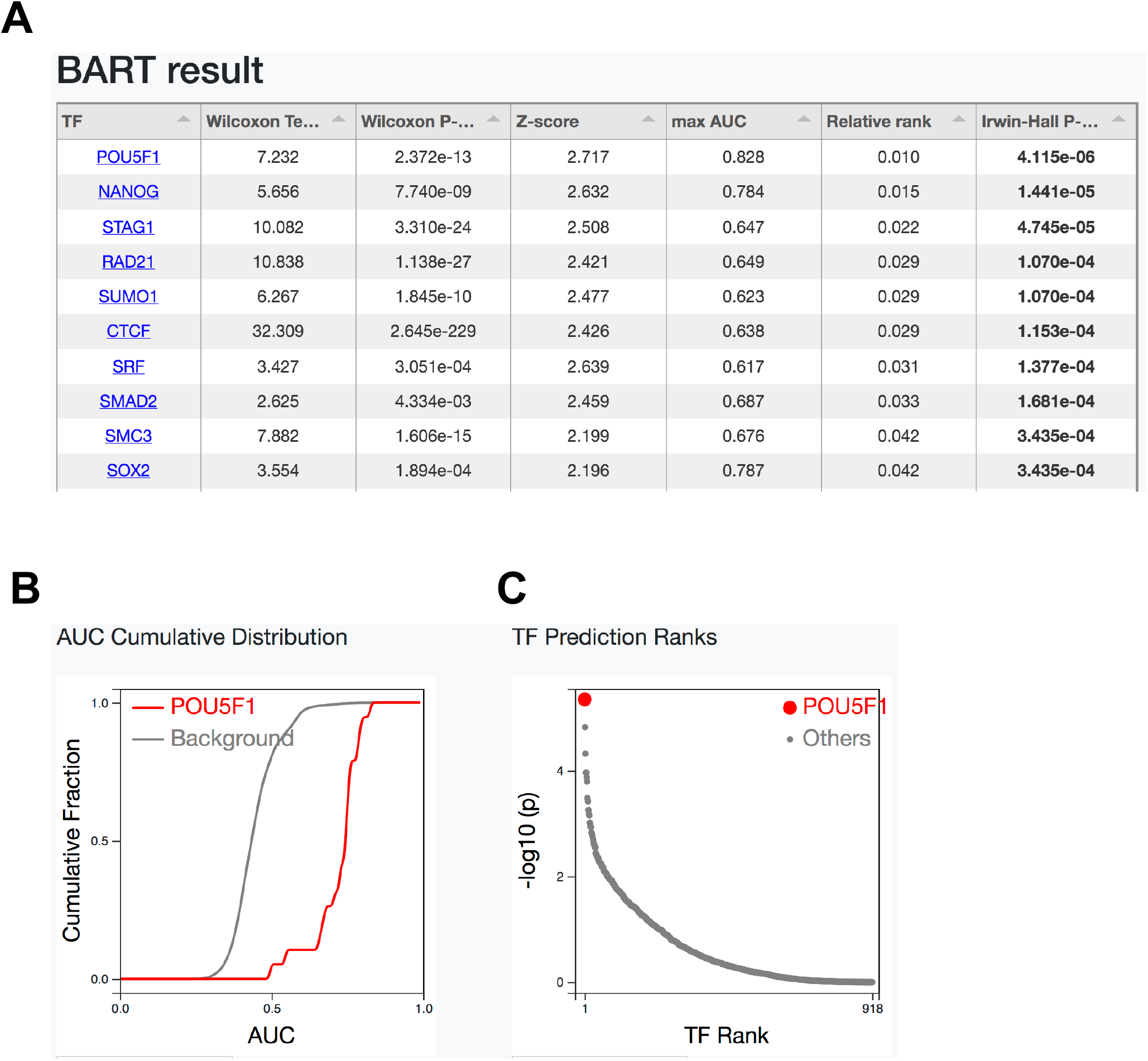
Example of BARTweb results. (**A**) Ranked list of identified TFs with quantification scores. (**B**) Cumulative distributions of association scores (AUC) of all ChIP-seq datasets for POU5F1 (red) compared with that of all other ChIP-seq datasets as background (grey). (**C**) Scatter plot of all TFs’ Irwin-Hall P-value score (−log_10_ P-value) against its rank. Selected TF (POU5F1) was labelled in red.

In the output TF table (Figure 3A), all available TFs (918 in human or 565 in mouse) will be displayed accompanied by six quantification scores in Columns 2-7. There are explanations on each column for definition reminder. Clicking on the column header can re-order the whole list by that score in a descending or ascending way.

- *Wilcoxon test statistic & P-value* (Columns 2 & 3): These two values indicate the level of association of each TF under the background of all other TFs. For each TF, we use Wilcoxon rank-sum test to compare the association scores from all ChIP-seq datasets for that TF with the association scores from all ChIP-seq datasets for other TFs.
- *Z-score* (Column 4): This value is to assess the specificity of each TF compared with a background model. We build background models using the Wilcoxon test statistics obtained from all annotated gene sets from the Molecular Signatures Database (MSigDB) (24) for gene set input or all H3K27ac ChIP-seq datasets from the data compendium for ChIP-seq read input, respectively.
- *Max AUC* (Column 5): The maximum association score among multiple ChIP-seq datasets of that TF.
- *Relative rank* (Column 6): The average rank of Wilcoxon test statistic, Z-score and maxAUC for each TF, divided by the total number of TFs.
- *Irwin-Hall P-value* (Column 7): This P-value indicates the integrative rank significance, using the Irwin-Hall distribution as the null distribution for unrelated ranks. The output TFs are ranked by this P-value by default. User can use 0.01 as a threshold for significant TF identification.

Example results shown in Figure 3 were authentically generated by BARTweb, using a gene set that were downregulated upon transcription factor OCT4 (POU5F1) was knocked down in a human embryonic stem cell line. This input gene set should include direct target genes of POU5F1. As expected, POU5F1 was identified as the top ranked regulator of the input gene set, whereas several other ESC-specific TFs such as NANOG and SOX2 were also identified.

Each TF in the table has a link to its corresponding analysis plots, including cumulative distribution of association scores (AUC) (Figure 3B) and rank-dot plot (Figure 3C). The cumulative distribution is plotted based on association scores of that TF comparing to all factors. When the curve of that TF appears on the right of all factors as background, it indicates that many ChIP-seq datasets for that TF have high association scores with the input genomic regulatory profile. The rank-dot plot is the most comprehensive and most straightforward plot in the analysis result. It plots Irwin-Hall P-value scores against absolute ranks of all TFs with the selected factor highlighted. Users can hover the mouse on other data points to find out which TF it is. The plots are in a high resolution of 600 dpi and can be downloaded and directly used in a manuscript.

Besides the interactive result panel, BARTweb also provides download links to all intermediate data files for further exploration, including selected H3K27ac samples from the adaptive lasso regression model, the genomic cis-regulatory profile and all TF ChIP-seq association scores. The regression information file tells which samples are selected along with regression coefficients and sample annotations including cell line, cell type or tissue type.

### BARTweb outperforms existing tools

To evaluate the performance of BARTweb on identifying the correct TFs that regulate the input gene set, we performed TF identification analysis using the gene sets derived from knockTF (25), a database of a comprehensive collection of a total of 570 human differential gene expression profiles with knockdown/knockout (KD/KO) of 308 TFs, and compared the BARTweb results with those generated from several other tools, including BART v1.1 (7), TFEA.ChIP (13), ChEA3 (14), Pscan (19), and HOMER (18). We summarized the features of the tools being compared in Supplementary Table S1. For each differential gene expression profile under a KD/KO TF, we used a fold-change cutoff of 1.5 to select the up- and down-regulated genes and conduct TF identification analyses separately. If the actual KD/KO TF was ranked among the top 10% of all TFs in the output and the corresponding P-value < 0.01 for either up- or down-regulated gene set, we declared that this tool yielded a true prediction on this dataset for this TF. Among the 570 differential expression datasets, 354 have their KD/KO TF included in the BARTweb TF library, and BARTweb had true predictions on 104 datasets, higher than the other tools (Figure 4A). If we focused on the number of unique TFs that each tool can successfully identify from the konckTF differentially expressed gene sets, BARTweb yielded true predictions for 61 TFs, also the highest among the tools tested (Figure 4B).

**Figure 4.**
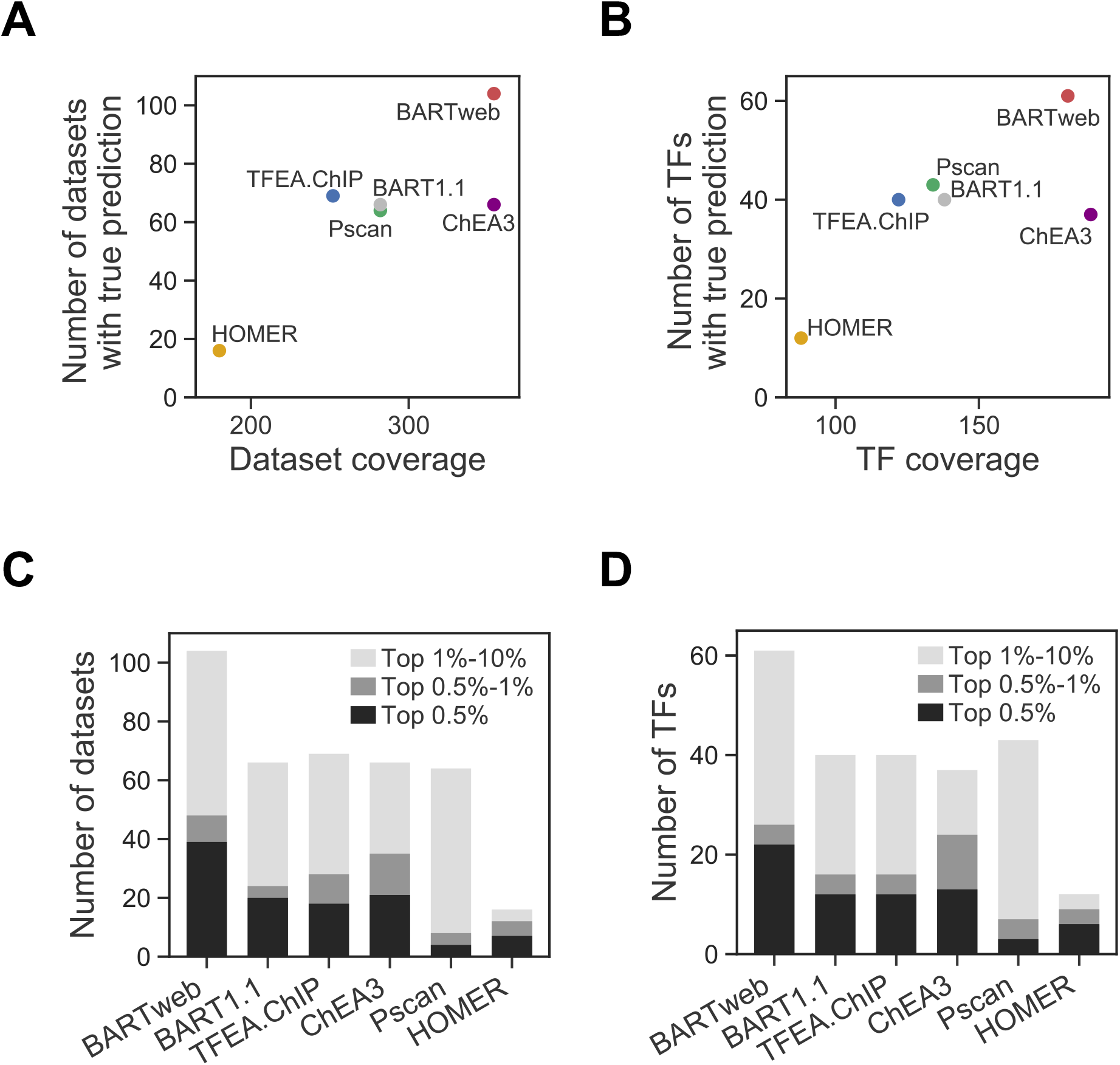
Performance comparison of BARTweb with 5 other tools on knockTF datasets. (**A**) Performance of each tool reflected by the number of knockTF datasets with true prediction (y-axis) against the number of knockTF datasets whose KD/KO TF were included in the tool (x-axis). (**B**) Performance of each tool reflected by the number of unique KD/KO TFs with true prediction (y-axis) against the number of unique KD/KO TFs included in the tool (x-axis). (**C**) Number of knockTF datasets with true prediction under different rank cutoffs for each tool. (**D**) Number of unique KD/KO TFs with true prediction under different rank cutoffs for each tool.

As general users tend to focus on only a few identified TFs ranked on top, the top 10% of all TFs could still be too many to follow up. Therefore, we further assessed the true prediction performances comparing the 5 tools, using top 0.5% and top 1% as cutoffs in addition to top 10%. In every case, we found that BARTweb still yielded true predictions on the highest number of gene expression datasets (Figure 4C) and the highest number of unique TFs (Figure 4D). In conclusion, we showed that BARTweb outperforms existing ChIP-seq-based and sequence motif-based tools in identifying regulatory TFs using the target gene sets from experimental data.

## DISCUSSION

We developed BARTweb, a web server for performing regulatory factor association analysis using a large collection of publicly available ChIP-seq datasets as the sole resource. This approach complements the commonly used sequence motif scan methods for TF identification, and has the unique advantage of utilizing in vivo protein-DNA interaction information across the genome for making biologically meaningful predictions. BARTweb has a user-friendly web interface, and can take in multiple types of input data. Despite integrating massive data with memory-intensive computing, BARTweb runs fairly fast by applying appropriate algorithms supported by statistical models.

Meanwhile, users should be aware of several limitations when using BARTweb for TF association analysis. First, as shown in Figure 4, the power of BARTweb is limited by the range of TFs that have existing ChIP-seq data from the public domain. The current data library of 918 factors might cover over half of human TFs, with the remaining hundreds of TFs still unable to be identified by BARTweb. While BARTweb is being maintained and updated, we expect that the TF coverage will keep growing, as we anticipate the publicly available ChIP-seq datasets will keep increasing. Secondly, similar to other tools, BARTweb does not consider cell-type specificity in the TF association analysis, although we do not think this is an issue. Because of the DNA sequence specificity of TF binding, genomic binding profiles of the same TF in different tissue/cell types are usually more similar to each other than binding profiles between different TFs in the same tissue/cell type (Supplementary Figure S2).

As a result, as long as the genomic cis-regulatory profile represents the binding profile of the functional TF, the BART algorithm is still able to find the correct TF, even from a different cell type, but unlikely to identify an irrelevant TF from a similar cell type. Nevertheless, as shown to have a superior performance than several similar tools, BARTweb is a powerful and easy-to-use bioinformatics web server for transcription factor analysis for different types of omics data. It can help biologists in gene regulation research interpret the experimental data and develop hypothesis for functional studies.

## SUPPLEMENTARY DATA

Supplementary Data include two supplementary figures, one supplementary table, and one supplementary algorithm.

## ACKNOWLEDGEMENTS

The authors would like to thank members of the Zang Laboratory for testing the web server, Byoung-Do Kim for support on research computing resources, and all users of the BARTweb beta version for helpful feedbacks.

---

**Supplementary Figure S1.**
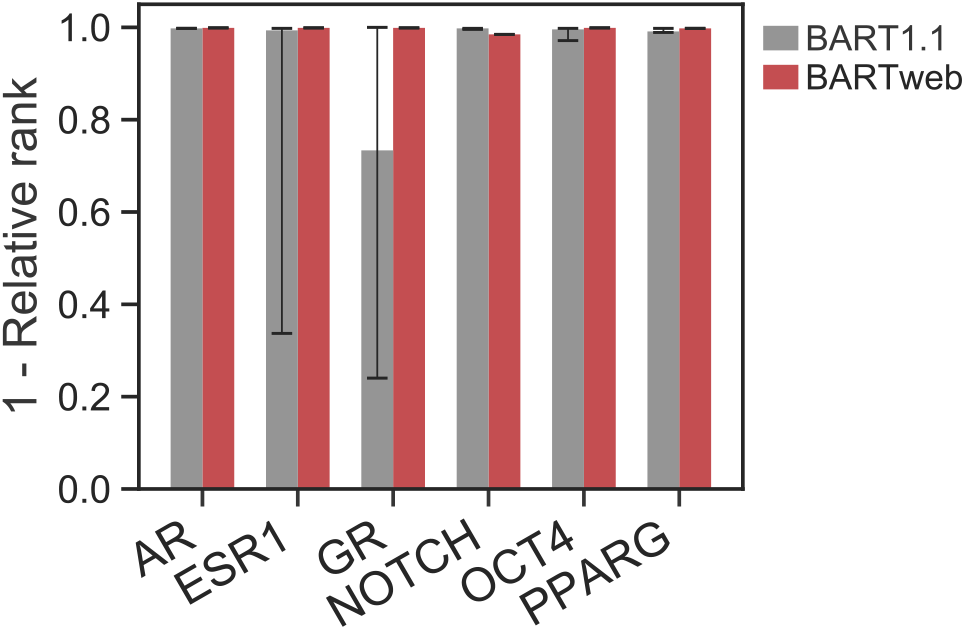
Bar plot comparing BARTweb and BART1.1 on functional TF identification results on six differentially expressed gene sets, collected from RNA-seq data upon activation of AR, ESR1, GR (NR3C1), NOTCH1, PPARG and knock-down of OCT4 (POU5F1). Each bar represents the median results from 20 runs; error bars represent the 25/75 percentiles.

**Supplementary Figure S2.**
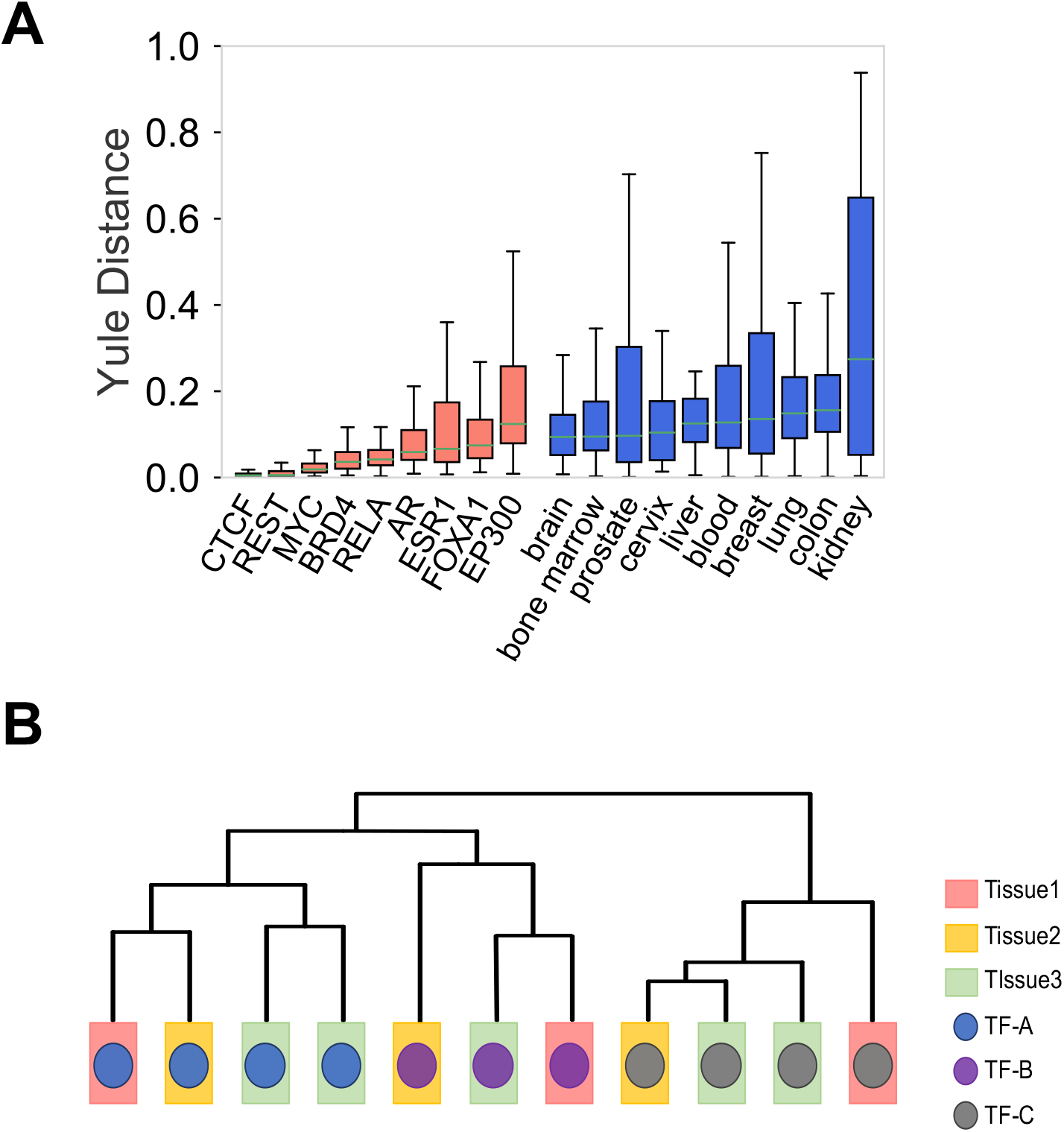
(**A**) Genome-wide TF binding specificity compared with tissue/cell-type specificity. For each TF with more than 50 ChIP-seq datasets among the 3485 collected human datasets, the Yule distance between each pair of datasets from different tissue types was calculated (orange). For each tissue type with more than ChIP-seq 50 datasets, the Yule distance between each pair of datasets of different factors was calculated (blue). Higher Yule distance indicates less similarity. P=0.0004, by Mann Whitney U test. (**B**) Schematic showing that TF binding patterns prevail over cell types in hierarchical clustering.

**Supplementary Table S1.**
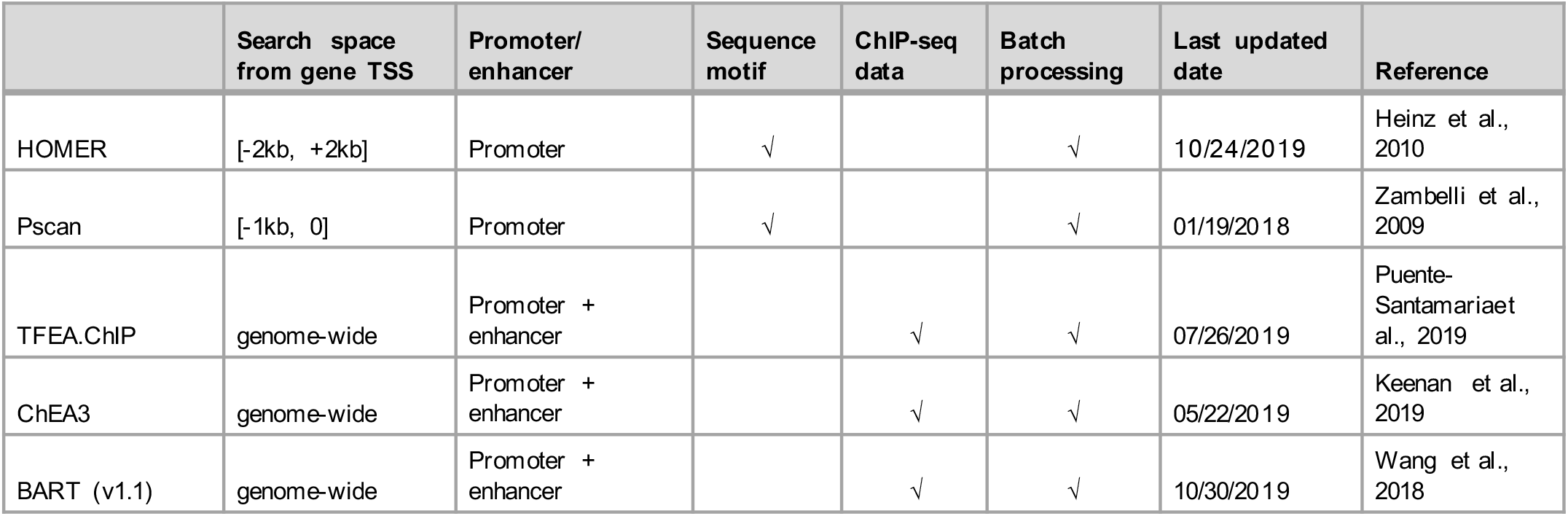
Existing tools being compared with BARTweb

Existing tools being compared with BARTweb
|  | Search space<br>from gene TSS | Promoter/<br>enhancer | Sequence<br>motif | ChIP-seq<br>data | Batch<br>processing | Last updated<br>date | Reference |
| --- | --- | --- | --- | --- | --- | --- | --- |
| HOMER | [-2kb, +2kb] | Promoter | √ |  | √ | 10/24/2019 | Heinz et al.,<br>2010 |
| Pscan | [-1kb, 0] | Promoter | √ |  | √ | 01/19/2018 | Zambelli et al.,<br>2009 |
| TFEA.ChIP | genome-wide | Promoter +<br>enhancer |  | √ | √ | 07/26/2019 | Puente-Santamaria et<br>al., 2019 |
| ChEA3 | genome-wide | Promoter +<br>enhancer |  | √ | √ | 05/22/2019 | Keenan et al.,<br>2019 |
| BART (v1.1) | genome-wide | Promoter +<br>enhancer |  | √ | √ | 10/30/2019 | Wang et al.,<br>2018 |

## Supplementary Algorithm 1 Algorithm

Logistic adaptive lasso

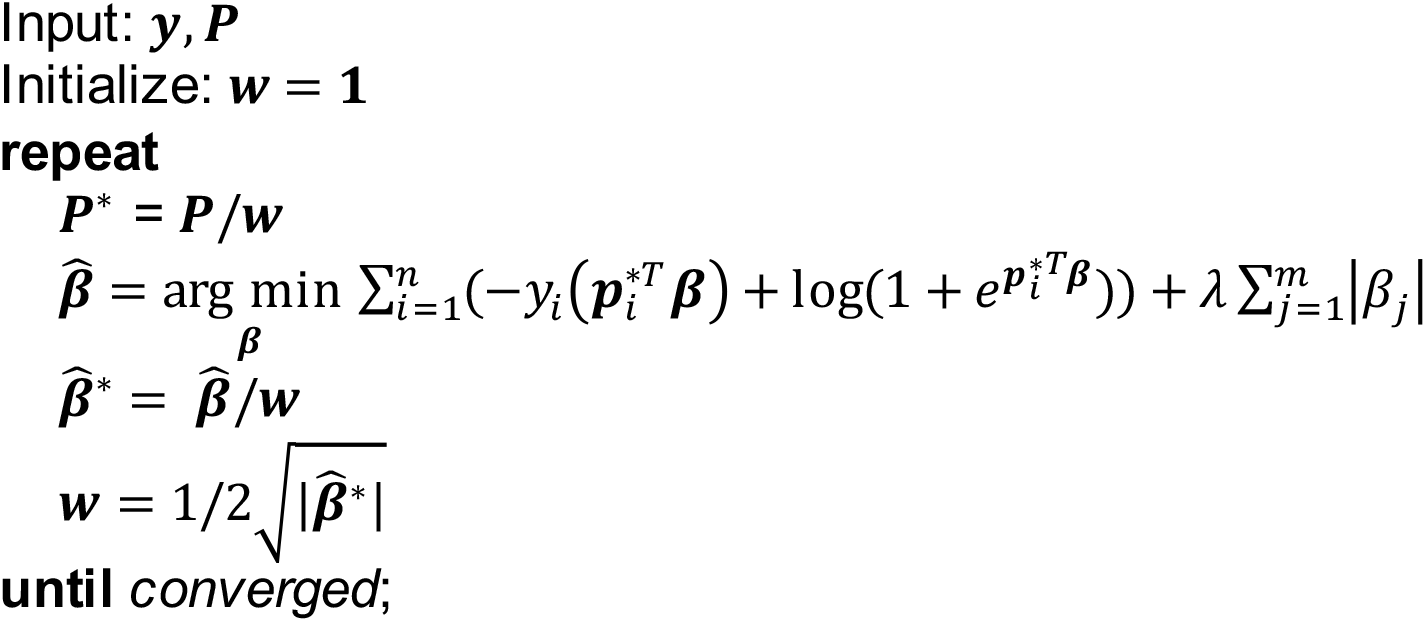

## REFERENCES

1. Lambert,S.A., Jolma,A., Campitelli,L.F., Das,P.K., Yin,Y., Albu,M., Chen,X., Taipale,J., Hughes,T.R. and Weirauch,M.T. (2018) The Human Transcription Factors. Cell, 172, 650–665. DOI: 10.1016/j.cell.2018.01.029.

2. Johnson,D.S., Mortazavi,A., Myers,R.M. and Wold,B. (2007) Genome-Wide Mapping of in Vivo Protein-DNA Interactions. Science, 316, 1497–1502. DOI: 10.1126/science.1141319.

3. Barski,A., Cuddapah,S., Cui,K., Roh,T.-Y., Schones,D.E., Wang,Z., Wei,G., Chepelev,I. and Zhao,K. (2007) High-Resolution Profiling of Histone Methylations in the Human Genome. Cell, 129, 823–837. DOI: 10.1016/j.cell.2007.05.009.

4. ENCODE Project Consortium (2012) An integrated encyclopedia of DNA elements in the human genome. Nature, 489, 57–74. DOI: 10.1038/nature11247.

5. Bernstein,B.E., Stamatoyannopoulos,J.A., Costello,J.F., Ren,B., Milosavljevic,A., Meissner,A., Kellis,M., Marra,M.A., Beaudet,A.L., Ecker,J.R., et al. (2010) The NIH Roadmap Epigenomics Mapping Consortium. Nature Biotechnology 2019, 28, 1045–1048. DOI: 10.1038/nbt1010-1045.

6. Mei,S., Qin,Q., Wu,Q., Sun,H., Zheng,R., Zang,C., Zhu,M., Wu,J., Shi,X., Taing,L., et al. (2017) Cistrome Data Browser: a data portal for ChIP-Seq and chromatin accessibility data in human and mouse. Nucleic Acids Res, 45, D658–D662. DOI: 10.1093/nar/gkw983.

7. Wang,Z., Civelek,M., Miller,C.L., Sheffield,N.C., Guertin,M.J. and Zang,C. (2018) BART: a transcription factor prediction tool with query gene sets or epigenomic profiles. Bioinformatics, 34, 2867–2869. DOI: 10.1093/bioinformatics/bty194.

8. Wang,S., Zang,C., Xiao,T., Fan,J., Mei,S., Qin,Q., Wu,Q., Li,X., Xu,K., He,H.H., et al. (2016) Modeling cis-regulation with a compendium of genome-wide histone H3K27ac profiles. Genome Res., 26, 1417–1429. DOI: 10.1101/gr.201574.115.

9. Parolia,A., Cieslik,M., Chu,S.-C., Xiao,L., Ouchi,T., Zhang,Y., Wang,X., Vats,P., Cao,X., Pitchiaya,S., et al. (2019) Distinct structural classes of activating FOXA1 alterations in advanced prostate cancer. Nature, 132, 3431–418. DOI: 10.1038/s41586-019-1347-4.

10. Shah,K.K., Whitaker,R.H., Busby,T., Hu,J., Shi,B., Wang,Z., Zang,C., Placzek,W.J. and Jiang,H. (2019) Specific inhibition of DPY30 activity by ASH2L-derived peptides suppresses blood cancer cell growth. Exp. Cell Res., 382, 111485. DOI: 10.1016/j.yexcr.2019.06.030.

11. Cheng,Q., Khoshdeli,M., Ferguson,B.S., Jabbari,K., Zang,C. and Parvin,B. (2019) YY1 is a Cis-regulator in the organoid models of high mammographic density. Bioinformatics, 10.1093/bioinformatics/btz812.

12. Jose,C.C., Wang,Z., Tanwar,V.S., Zhang,X., Zang,C. and Cuddapah,S. (2019) Nickel-induced transcriptional changes persist post exposure through epigenetic reprogramming. Epigenetics Chromatin, 12, 75–15. DOI: 10.1186/s13072-019-0324-3.

13. Puente-Santamaria,L., Wasserman,W.W. and Del Peso,L. (2019) TFEA.ChIP: A tool kit for transcription factor binding site enrichment analysis capitalizing on ChIP-seq datasets. Bioinformatics, 29, 1922. DOI: 10.1093/bioinformatics/btz573.

14. Keenan,A.B., Torre,D., Lachmann,A., Leong,A.K., Wojciechowicz,M.L., Utti,V., Jagodnik,K.M., Kropiwnicki,E., Wang,Z. and Ma’ayan,A. (2019) ChEA3: transcription factor enrichment analysis by orthogonal omics integration. Nucleic Acids Res, 47, W212–W224. DOI: 10.1093/nar/gkz446.

15. Subramanian,A., Tamayo,P., Mootha,V.K., Mukherjee,S., Ebert,B.L., Gillette,M.A., Paulovich,A., Pomeroy,S.L., Golub,T.R., Lander,E.S., et al. (2005) Gene set enrichment analysis: a knowledge-based approach for interpreting genome-wide expression profiles. PNAS, 102, 15545–15550. DOI: 10.1073/pnas.0506580102.

16. Chèneby,J., Gheorghe,M., Artufel,M., Mathelier,A. and Ballester,B. (2018) ReMap 2018: an updated atlas of regulatory regions from an integrative analysis of DNA-binding ChIP-seq experiments. Nucleic Acids Res, 46, D267–D275. DOI: 10.1093/nar/gkx1092.

17. Kuleshov,M.V., Jones,M.R., Rouillard,A.D., Fernandez,N.F., Duan,Q., Wang,Z., Koplev,S., Jenkins,S.L., Jagodnik,K.M., Lachmann,A., et al. (2016) Enrichr: a comprehensive gene set enrichment analysis web server 2016 update. Nucleic Acids Res, 44, W90–7. DOI: 10.1093/nar/gkw377.

18. Heinz,S., Benner,C., Spann,N., Bertolino,E., Lin,Y.C., Laslo,P., Cheng,J.X., Murre,C., Singh,H. and Glass,C.K. (2010) Simple combinations of lineage-determining transcription factors prime cis-regulatory elements required for macrophage and B cell identities. Mol. Cell, 38, 576–589. DOI: 10.1016/j.molcel.2010.05.004.

19. Zambelli,F., Pesole,G. and Pavesi,G. (2009) Pscan: finding over-represented transcription factor binding site motifs in sequences from co-regulated or co-expressed genes. Nucleic Acids Res, 37, W247–W252. DOI: 10.1093/nar/gkp464.

20. Zou,H. (2006) The Adaptive Lasso and Its Oracle Properties. Journal of the American Statistical Association, 101, 1418–1429. DOI: 10.1198/016214506000000735.

21. Hesterberg,T., Choi,N.H., Meier,L. and Fraley,C. (2008) Least angle and ℓ1 penalized regression: A review. Statistics Surveys, 2, 61–93. DOI: 10.1214/08-SS035.

22. Efron,B., Hastie,T., Johnstone,I. and Tibshirani,R. (2004) Least angle regression. The Annals of Statistics, 32, 407–499. DOI: 10.1214/009053604000000067.

23. Zheng,R., Wan,C., Mei,S., Qin,Q., Wu,Q., Sun,H., Chen,C.-H., Brown,M., Zhang,X., Meyer,C.A., et al. (2019) Cistrome Data Browser: expanded datasets and new tools for gene regulatory analysis. Nucleic Acids Res, 47, D729–D735. DOI: 10.1093/nar/gky1094.

24. Liberzon,A., Birger,C., Thorvaldsdóttir,H., Ghandi,M., Mesirov,J.P. and Tamayo,P. (2015) The Molecular Signatures Database (MSigDB) hallmark gene set collection. Cell Systems, 1, 417–425. DOI: 10.1016/j.cels.2015.12.004.

25. Feng,C., Song,C., Liu,Y., Qian,F., Gao,Y., Ning,Z., Wang,Q., Jiang,Y., Li,Y., Li,M., et al. (2019) KnockTF: a comprehensive human gene expression profile database with knockdown/knockout of transcription factors. Nucleic Acids Res, 13, 613. DOI: 10.1093/nar/gkz881.

